# Conserved accessory proteins encoded with archaeal and bacterial Type III CRISPR-Cas gene cassettes that may specifically modulate, complement or extend interference activity

**DOI:** 10.1101/262675

**Authors:** Shiraz A. Shah, Omer S. Alkhnbashi, Juliane Behler, Wenyuan Han, Qunxin She, Wolfgang R. Hess, Roger A. Garrett, Rolf Backofen

## Abstract

A study was undertaken to identify conserved proteins that are encoded either within, or directly adjacent to, *cas* gene cassettes of Type III CRISPR-Cas interference modules. These Type III modules are especially versatile functionally and have been shown to target and degrade dsDNA, ssDNA and ssRNA. In addition, the interference gene cassettes are frequently intertwined with other accessory genes, including genes encoding CARF domains, some of which are likely to be cofunctional. Using a comparative genomics approach, and defining a Type III association score accounting for coevolution and specificity of flanking genes, we identified and classified 39 new Type III associated gene families. Most archaeal and bacterial Type III modules were seen to be flanked by several accessory genes, around half of which did not encode CARF domains and remain of unknown function. Non-CARF accessory genes were found to be more diverse than their CARF counterparts, encoding nuclease, helicase, protease, ATPase, transporter and transmembrane domains and including a considerable fraction that encoded no known domains. The diversity of non-CARF Type III accessory genes found in this study suggests that additional families exist which remain undetected because of the limited number of annotated genomes currently available. The method employed is scalable for potential application on metagenomic data once automated pipelines for annotation of CRISPR-Cas systems have been developed. All accessory genes found in this study are presented online in a readily accessible and searchable format for researchers to audit their model organism of choice: http://accessory.crispr.dk.

## Introduction

Type III CRISPR-Cas systems have been classified into four main subtypes A to D of which subtypes III-A and III-B have been studied extensively (Vestergaard et al. 2014; Makarova et al. 2015). They often coexist within cells carrying Type I systems and are assumed to complement the latter functionally. However, whereas Type I systems specifically target dsDNA, Type III systems can degrade dsDNA, ssDNA and ssRNA (Marraffini and Sontheimer 2008; Hale et al. 2009; Peng et al. 2015; Elmore et al. 2016; Zhang et al. 2016; Han et al. 2017a; Tamulaitis et al. 2017). Importantly, an earlier survey of archaeal *cas* gene cassettes revealed that Type III modules are exceptional in that they frequently carry accessory genes either within or immediately bordering their core *cas* gene modules (Vestergaard et al. 2014). Some of these accessory genes encode putative proteases or nucleases suggesting that they provide additional functions that modulate, complement or extend, Type III interference functions.

Some accessory *cas* genes were initially identified during the first comprehensive CRISPR-Cas subtype classification (Haft et al. 2005). Core *cas* genes were found to be conserved across most CRISPR-Cas subtypes, while a different class of *cas* genes displayed a more variable distribution and were not consistently associated with specific subtypes. Such genes were designated csx and six gene families, *csx1-6*, were classified (Haft et al. 2005). In the next major CRISPR-Cas systematics update, some accessory *cas* gene families were merged into the existing, and new, core *cas* gene families, with only the *csx1* and *csx3* families remaining separate (Makarova et al. 2011). This more robust definition of core *cas* gene families enabled a more rigorous distinction to be made from eventual new accessory genes. Building on this, an archaea-specific study identified almost twenty new *cas* accessory gene families that were considered distinct from core *cas* gene families, an inference that was reinforced by their variable distribution amongst CRISPR-Cas subtypes (Vestergaard et al. 2014). Most of the accessory genes were found to be associated with Type III systems, and most of the encoded proteins were related to the original Csx1 family proteins (Haft et al. 2005). Two additional Type III subtypes, III-C and III-D were also introduced, and they were seen to be associated with many of the same accessory genes as those linked to Type III-A and III-B modules (Vestergaard et al., 2014). Later, the Csx1 protein family was formally defined by its ligand binding CARF domain and was annotated in both archaea and bacteria (Makarova et al. 2014). In addition, a few bacterial accessory proteins families lacking CARF domains were found and added to the TIGRFAMs database by the author of the original study (Haft et al. 2005). The latest CRISPR-Cas systematics update (Makarova et al. 2015) adopted the new Type III subtypes and integrated many of the newly identified archaeal and bacterial accessory genes families. Moreover, subtype III-D was extended to include disparate bacterial Type III systems that had previously defied classification. As a result, their core genes were relatively poorly characterised and were sometimes annotated as accessory genes. Nevertheless, Makarova et al. (2015) provided detailed annotations of all known archaeal and bacterial CRISPR-Cas systems as Supplementary Material for future reference; the present study is based on those data.

Although experimental studies on non-CARF accessory proteins have been limited to Cmr7 from *Sulfolobus* (Zhang et al. 2012), numerous studies have investigated the role of common CARF family accessory proteins such as Csx1, Csm6 and Csa3. The crystal structure of Csa3 was first resolved in 2011 (Lintner et al. 2011) and it demonstrated that the C-terminal DNA binding HTH domain was likely to be under allosteric control of the N-terminal Rossman fold domain implicated in dinucleotide ligand sensing, of the type that is normally involved in signal recognition. Furthermore, a crystal structure of Cmr2, with its cyclase domain, was also compatible with the occurrence of such a signal (Zhu and Ye 2012). These early insights were later undermined by genetic studies on Type III systems (Deng et al. 2013; Hatoum-Aslan et al. 2014) that linked the presence of Csx1 and Csm6 to a DNA interference phenotype. This was enigmatic at the time because several crystallographic, biochemical and bioinformatic studies had predicted that the proteins must be primarily RNases (Makarova et al. 2012; Kim et al. 2013; Niewoehner and Jinek 2016). A key proposal was subsequently made that Csx3 was a distant CARF family member and that the ligands sensed by CARF proteins in general were cyclic oligonucleotides (Yan et al. 2015); (Topuzlu and Lawrence 2016). This hypothesis was confirmed in three recent independent studies (Kazlauskiene et al. 2017; Niewoehner et al. 2017; Han et al. 2017b), two of which also showed that the cyclic oligoadenylate (cOA) signaling molecules were synthesised by the polymerase domain of Cas10 upon recognition of its target (Makarova et al. 2006; Zhu and Ye 2012). Apparently any target recognition by cellular Type III interference complexes triggers cOA synthesis which, in turn, ensures the coordinated activation of a potentially complex and layered defence response that is dependent on intracellular CARF proteins.

Although CARF proteins constitute the most widespread Type III accessory proteins, accessory proteins lacking CARF domains are much more diverse and almost as widespread. For example, it was shown that archaeal Type III interference gene cassettes are often flanked by diverse accessory genes encoding protein domains associated with nucleases, proteases, helicases, toxins or transcriptional regulators (Zhang et al. 2012; Deng et al. 2013; Vestergaard et al. 2014). A search has already been undertaken specifically for CARF-encoding genes in both archaea and bacteria, and it yielded a comprehensive catalogue of CARF domain-carrying proteins (Makarova et al. 2014). Here we identify new families of non-CARF *cas* accessory genes associated with Type III modules, in addition to those encoding CARF domains, in archaeal and bacterial genomes. Since non-CARF accessory genes do not share a common sequence motif or domain, they are particularly challenging to identify. Our method relies on a guilt-by-association approach, coupled with criteria for coevolution and specificity to exclude false positives. Type I, II and IV *cas* gene cassettes were not examined systematically because the incidence of accessory genes is too low to distinguish signal from noise with our method given the limited number of annotated genomes available, although an independent study has been undertaken for these CRISPR-Cas types (Shmakov et al., 2018). In the present study, by focusing on Type III systems, we are better able to identify reliable candidate accessory genes from a spurious gene background.

## Results

From a total of 1263 archaeal and bacterial genomes in which CRISPR-Cas cassettes are fully annotated (Makarova et al. 2015), 381 genomes encoded 512 distinct Type III genetic modules which were surveyed for accessory genes (Table 1). Selecting five genes immediately upstream and downstream from each module yielded a total of 4467 genes. This number was less than the theoretical total of 5120 genes because neighbouring adaptation modules, Type I interference modules and *cas*6 genes were omitted from the analysis. All the encoded protein sequences were aligned with one another and a custom similarity metric (Vestergaard et al. 2014) was calculated and used for Markov clustering (Enright et al. 2002). This yielded 231 gene clusters each with more than three members. Cas association scores were calculated for each gene cluster using information about Type III coevolution, Type III subtype specificity, and host genome self-similarity. Gene clusters with low Cas association scores were considered spurious and disregarded. The Cas association score ranged from 100 to −100 and a cut off of 24 was set by comparing the results to the manually curated accessory gene families obtained in the previous archaeal genome study (Vestergaard et al. 2014). Thereafter, 76 of the 231 Type III-associated gene clusters were considered significant; the remainder were discarded. The 76 gene clusters encompassed a total of 982 potential accessory genes distributed across the 512 Type III modules. This translated, on average, to two accessory genes per Type III module, varying from no additional genes to more than the core Type III genes (Figure 1). Properties of the 76 gene families are summarised (Table 2) and a more elaborate version (Table S1) is available online.

**Figure 1:**
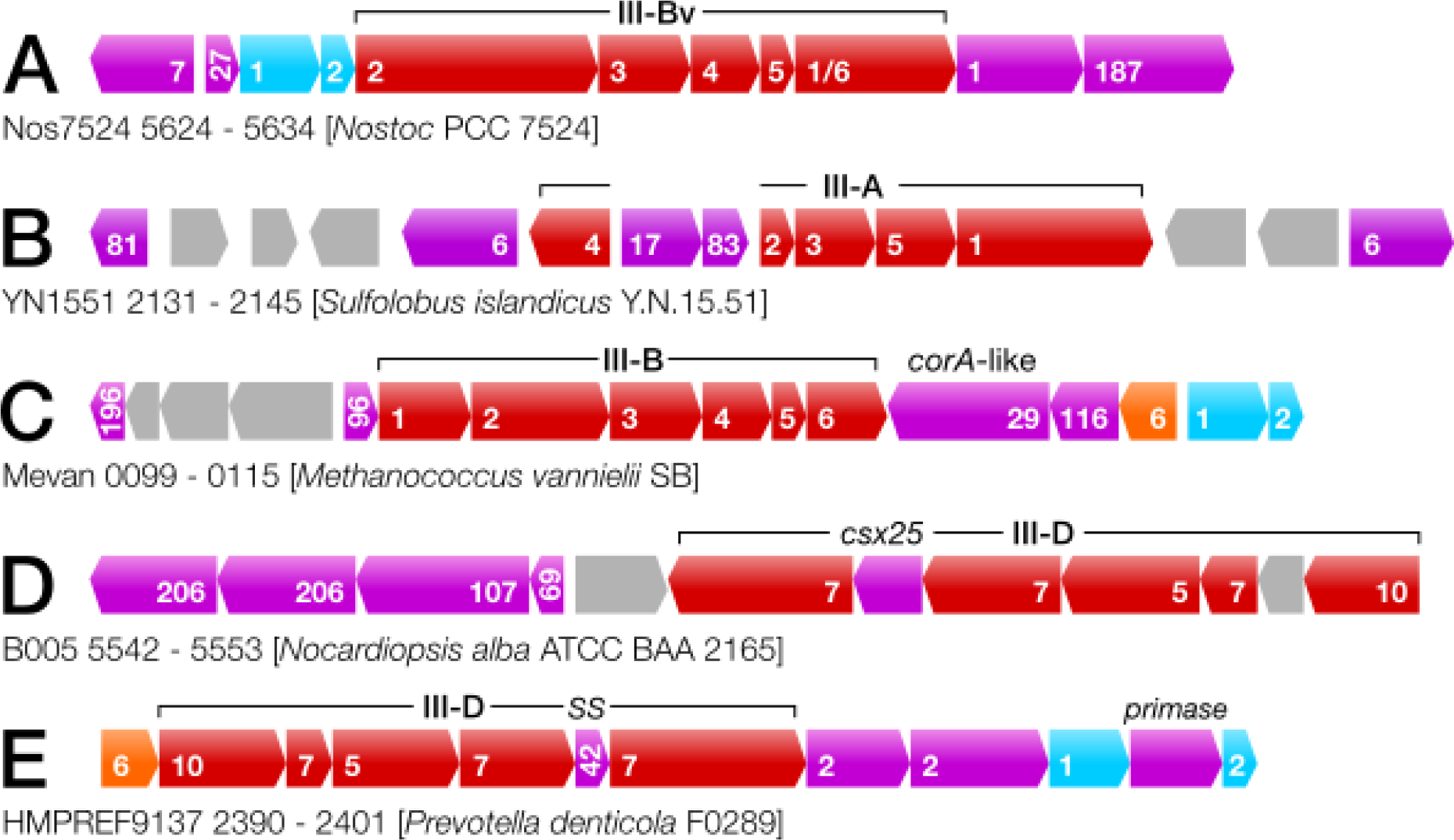
Gene maps of example Type III gene cassettes including accessory genes. Core Type III genes are drawn in red with *csm*/*cmr* numbers for Type III-A/B modules, and *cas* numbers for Type III-D modules. *cas6* and the adaptation module genes are orange and bIue respectively, also with *cas* gene numbers. Accessory genes are drawn in purpIe with the gene cluster number indicated.

**Table 1:** Summary of results from the current study compared to previous studies that identified accessory *cas* genes, divided between archaea and bacteria. The previous studies were limited to a (a) archaeal genomes (Vestergaard et al. 2014) and (b) a representative subset of genomes (Makarova et al. 2014). The present study is more comprehensive.

|  | present study |  | Makarova 2014 |  | Vestergaard 2014 |
| --- | --- | --- | --- | --- | --- |
|  | Ar | Ba | Ar | Ba | Ar |
| Number of genomes surveyed | 217 | 2534 | 172 | 484 | 159 |
| Number of genomes with CRISPR-Cas systems | 131 | 1131 | 50 | 133 | 124 |
| Number of genomes with Type III systems | 78 | 304 | 34 | 114 | 83 |
| Total number of Type III systems | 110 | 402 | 61 | 251 | 125 |
| Number of genomes with accessory genes | 67 | 262 | 78 | 180 | 76 |
| Total number of putative accessory genes | 248 | 734 | 190 | 454 | 239 |
| Accessory genes found outside Cas cassettes | 0 | 0 | 104 | 152 | 0 |
| Accessory genes associated to Type III systems | 248 | 734 | 61 | 251 | 210 |
| Total number of non-CARF accessory genes found | 136 | 369 | 0 | 0 | 135 |

**Table 2:** List of putative accessory gene clusters conserved near Type III genetic modules which passed the Type III association score cut-off (>24). A few gene families with lower scores are also included because they have been confirmed as accessory proteins in earlier studies. For each putative accessory protein family, the cluster id, the size (i.e. number of members per cluster), and the calculated Type III association score are listed. An example (gene-) locus id is also provided for reference. Names are provided for accessory protein families identified in earlier studies (Haft et al. 2005; Vestergaard et al. 2014; Makarova et al. 2014). Thirty nine of 76 putative accessory protein families are newly identified. Commonly associated Type III subtypes are listed for each putative accessory protein family. Unclassified variant subtypes are indicated by ‘III’. Most accessory gene families can function with different Type III subtypes. A predicted function is given in the last column.

| cluster | size | score | example | known | subtype | annotation |
| --- | --- | --- | --- | --- | --- | --- |
| 1 | 116 | 73.88 | JTY_2831 | Csx1/Csm6 | A,B,D | CARF+HEPN |
| 2 | 94 | 69.25 | Selin_1039 | Csx1 | A,B,D | CARF+HEPN |
| 3 | 69 | 62.33 | FFONT_0074 | Csm6 | A,B,D | CARF+RelE |
| 5 | 64 | 59.36 | ERE_12150 | Csx19/24 | (A,B),D | core gene (SS) |
| 6 | 61 | 58.51 | TOPB45_1115 | Csx1 | A,B,(C,D) | CARF+HEPN |
| 7 | 51 | 42.59 | Tph_c24580 | WYL/Csx1 | A,B,D | CARF+WYL |
| 9 | 49 | 37.91 | Vdis_1148 | Csx1 | (A),B,(C)D | CARF+Nuclease |
| 11 | 41 | 35.59 | Thebr_0941 | Csx15/20 | A,B,D | peptidase |
| 17 | 31 | 24.7 | VMUT_1481 | Cas_RecF | A,B,(C,D) | SMC ATPase |
| 23 | 20 | 4.61 | CYB_0598 | Csx1 | (A),B,D | found elsewhere |
| 25 | 18 | 55.83 | Dtur_0613 | Csx3 | A,B,D | Csx3 (CARF) |
| 27 | 17 | 50.07 | Cyan10605_3521 | Csx18 | B,(D) | adaptation associated |
| 28 | 17 | 66.76 | CYB_0586 | Csx21 | 17 | Type III-D associated |
| 29 | 14 | 45.91 | CBO2177 | CorA | B,(C),D | CorA-like |
| 33 | 13 | 10.47 | M1425_0883 | Csa3 | (A),B,C | Type I-A associated |
| 35 | 12 | 61.07 | CYB_0599 | Csx3 | (A),B,D | Csx3-AAA |
| 36 | 11 | 50.71 | Nos7107_4284 | WYL | (A,B),D | HTH-WYL |
| 37 | 11 | 56.62 | SacN8_09300 | Csx26 | III,A,(B) | HNH nuclease |
| 38 | 10 | 54.35 | Mefer_0963 | **new** | A,B | unknown |
| 42 | 9 | 74.4 | CFT03427_1632 | Csx19 | D | core gene (SS) |
| 43 | 9 | 58.83 | Ferpe_1557 | **new** | A,B | Lon protease |
| 45 | 9 | 28.54 | Marme_0670 | PD-DExK | A,B,D | possible nuclease |
| 47 | 9 | 27.09 | SacN8_09290 | protease | III,A,B,D | peptidase |
| 50 | 8 | 61.07 | Swol_2530 | Csx13 | (A),B,D | TM+CARF |
| 55 | 7 | 44.44 | Nos7107_2836 | **new** | B,D | ABC permease |
| 57 | 7 | 20.46 | Calkr_2542 | HerA | A,B,(C) | Type III coevolved |
| 58 | 7 | 33.01 | VMUT_1471 | NurA | A,B | Type III coevolved |
| 59 | 7 | 55.84 | Nos7107_2826 | **new** | A,B,D | trans-membrane (TM) |
| 64 | 7 | 43.07 | SacRon12I_09300 | **new** | III,A | unknown |
| 67 | 6 | 40.03 | Caur_2269 | **new** | (A),B | AAA-Csx3 |
| 69 | 6 | 29.69 | B005_5545 | **new** | D | unknown |
| 76 | 6 | 39 | LS215_0703 | **new** | III,A,D | unknown |
| 77 | 6 | 29.73 | NE0116 | Csx16 | A,B,D | a.k.a. cas_VVA1548 |
| 80 | 6 | 27.48 | PTH_0706 | **new** | B,C,D | AAA+ ATPase |
| 81 | 6 | -9.48 | YN1551_2131 | Csx1 | (A,B),D | Type I associated |
| 83 | 6 | 88.69 | VMUT_1493 | cas_RFas | A,(B) | cluster 17 associated |
| 84 | 6 | 66.57 | TTX_1229 | cas_RFas | A,(B) | cluster 17 associated |
| 87 | 6 | 74.65 | SSO1986 | Cmr7 | B | Cmr7 |
| 93 | 5 | 30.57 | Thebr_0950 | **new** | B,C,D | poss. AAA ATPase |
| 96 | 5 | 42.25 | Mvol_0529 | Mvol_0529-fam | B | DNA binding C-ter. |
| 104 | 5 | 39.74 | Msed_1167 | Csx1 | A,B,D | CARF+PIN |
| 107 | 5 | 84.37 | B005_5544 | Csx1 | B,D | CARF |
| 108 | 5 | 78.8 | Pcal_0278 | cas_RFas | B | cluster 17 associated |
| 116 | 4 | 57.92 | Rru_A0181 | **new** | B,D | cluster 29 associated |
| 121 | 4 | 62.04 | Desac_1715 | Csx23 | A,B,D | unknown |
| 123 | 4 | 37.09 | Caur_2303 | **new** | B,D | unknown |
| 124 | 4 | 44.42 | PCC7418_1341 | **new** | B | unknown |
| 139 | 4 | 28.91 | Tthe_0931 | **new** | A,B,D | HKD+Snf2 |
| 146 | 4 | 68.43 | SSO1421 | **new** | D | DNA binding HTH |
| 152 | 3 | 25.85 | Cylst_6373 | **new** | C | Type III-C specific |
| 156 | 3 | 28.1 | Metin_0159 | **new** | B,C | oxidoreductase |
| 159 | 3 | 27.06 | Calkr_2554 | **new** | A,B,C | methyl transferase |
| 162 | 3 | 40.43 | SYO3AOP1_0653 | **new** | III,A,B | unknown |
| 166 | 3 | 32.1 | Hbut_0719 | **new** | B,D | poss. crRNA proc. |
| 168 | 3 | 31.1 | slr7083 | **new** | B | unknown |
| 173 | 3 | 70 | Adeg_0988 | Csx21 | III,B,D | unknown |
| 174 | 3 | 74.3 | Adeg_0809 | **new** | B,C | nucleotidyl trans. |
| 178 | 3 | 37.97 | Csac_0071 | **new** | A,B,D | putative invertase |
| 181 | 3 | 25.55 | Tpen_1359 | **new** | A,B,C | Diadenylate cyclase |
| 186 | 3 | 28.77 | RoseRS_2594 | **new** | A,C,D | C-3',4' desaturase |
| 187 | 3 | 39 | Dole_0738 | **new** | A,B | NERD+UvrD |
| 189 | 3 | 50.73 | Hoch_5581 | **new** | B,D | kinase |
| 193 | 3 | 43.78 | Thebr_0949 | **new** | B,D | poss. alt. Cas3 |
| 194 | 3 | 26.83 | Mcup_1132 | **new** | B,D | AA transporter |
| 196 | 3 | 46.29 | TREPR_1099 | **new** | III,A,B | unknown |
| 198 | 3 | 40.67 | Mrub_1490 | **new** | B | TPR protein |
| 205 | 3 | 67.5 | MLP_11360 | **new** | D | unknown |
| 206 | 3 | 29.39 | Ndas_2980 | **new** | D | TAP-like protein |
| 211 | 3 | 51.67 | Pars_1112 | cas_RFas | B | unknown |
| 212 | 3 | 39.67 | RoseRS_0371 | **new** | C,D | SNc+LTD |
| 214 | 3 | 30.16 | Rcas_4246 | **new** | B,C | GlgB |
| 216 | 3 | 25.6 | SSA_1254 | **new** | A,B | adaptation associated |
| 222 | 3 | 44 | Vpar_1800 | **new** | A,D | unknown |
| 223 | 3 | 43.33 | YG5714_0635 | **new** | III,D | unknown |
| 227 | 3 | 36.73 | Sfum_1354 | PrimPol | A,B | adaptation polymerase |
| 230 | 3 | 28.76 | TVNIR_1454 | **new** | B | RecX family protein |

### Online availability of results

The complete set of results is provided online in an indexed and readily searchable format for further validation(Table S1, http://accessory.crispr.dk). Multiple sequence alignments are provided for protein sequences encoded by each gene cluster. Gene names, and host genome accession numbers are given for each gene together with the adjacent Type III module subtype (Table S2, http://accessory.crispr.dk/genes.html). Profile-profile matches to the sequence family databases Pfam (Punta et al. 2012), TIGRFAMs (Haft et al. 2005) and CDD (Makarova et al. 2011) are also presented for each gene family, along with matches to protein structures in PDB (Burley et al. 2017). Finally, gene maps have been drawn for each of the 982 separate putative accessory genes found, illustrating their genomic context including the neighbouring Type III genetic module and other accessory genes or core *cas* genes from other systems flanking the same module. Clicking the gene maps links to the gene entry in the RefSeq database, enabling easy retrieval of protein or gene sequences, along with the option of initiating a BLAST or CDD search with an additional click.

### Largest gene clusters

Of the five largest gene clusters (1 to 5), the first three had 116, 94 and 69 members, respectively (Table 2). All gave highly significant profile-profile matches to known CARF protein profiles (e.g. TIGR02710 or Csx1 and the 5FSH PDB entry for the Csm6 crystal structure from *Thermus thermophilus*). This result confirms the earlier observation for archaea that CARF domain-encoding genes comprise the most dominant class of accessory genes. The largest cluster with no CARF match, cluster 4, had a low Cas association score and was considered spurious and not included in Table 2. This particular gene cluster encodes an ancient, ubiquitous, family of ABC-transporters with high sequence conservation that would probably appear enriched regardless of the genomic locus under survey. Gene cluster 5 with 64 members was almost exclusively associated with the less well characterised III-D subtype modules. These genes were always closely linked with their cognate Type III genetic modules and co-transcribed with the core *cas* genes. The genes encode a relatively small protein (about 160 aa) with no significant matches in sequence databases. Therefore, we inferred that the gene encodes the small subunit (SS), or Cmr5 analog, of an uncharacterised subfamily of Type III-D modules (Figure 1E) and that cluster 5 represents a core Type III-D gene family rather than an accessory gene family.

### Most significantly associated gene clusters

The five most significant accessory gene clusters according to the calculated Cas association score were clusters 83, 107, 108, 87 and 42 (Table 2). Cluster 83 carries six genes all located adjacent to Type III-A gene cassettes from crenarchaeal orders including the thermoacidophilic Sulfolobales and the thermoneutrophilic Thermoproteales. These genes are invariably located immediately adjacent to cluster 17 genes (Figure 1B) encoding a putative ATPase domain protein (330 aa) (Table 2) common to DNA repair proteins of the SMC (Structural Maintenance of Chromosomes) type including rad50 and recF. The protein sequences that make up cluster 83 average 160 aa and yield no good profile matches in public databases. The cluster 83-17 pair of accessory genes is also accompanied by gene clusters 6 and 84 (Figure 1B), the former of which encodes a CARF protein of the MJ1666 type (Haft et al. 2005), while genes of the latter are similar in size to those of cluster 83 and show weak sequence similarity.

Cluster 107 has five gene members present in divergent bacterial genera *Nocardiopsis* and *Thermus*, members of the Actinobacterial and Thermus-Deinococcus phyla, respectively. Cluster 107 proteins produce significant sequence matches to known CARF domain-containing proteins but are larger (about 700 aa) and long regions in the middle and at the C-terminal end yield no profile-profile matches in databases. Thus, while it is likely that these CARF proteins are also activated by cOA, their effector function remains obscure. In *Nocardiopsis* the host carries subtype III-D modules (Figure 1D) and also harbours cluster 206 genes which encode a protease of the AB hydrolase family. Typically for CARF proteins, cluster 107 is found associated with different subtypes III-B and III-D.

Cluster 108 genes are found amongst crenarchaeal thermoneutrophiles, often in multiple divergent copies immediately adjacent to cluster 17 genes, similar to the coexistence of genes of clusters 17, 83 and 84. Cluster 108 proteins (about 180 aa) show no sequence similarity to cluster 83/84 proteins and yield no good matches to protein family databases. However, they contain a transmembrane (TM) signal at the N-terminal end consistent with the protein being membrane bound.

Cluster 87 corresponds to *cmr7* (Zhang et al. 2012) and is an additional *cas* gene so far exclusive to *Sulfolobus* species. Exceptionally Cmr7 has been characterised experimentally and shown to associate tightly with the Cas10/Cmr3 sub-complex within the cognate Type III-B effector complex (Zhang et al. 2012). Cluster 87 genes are associated exclusively with the Cmr-b subclass of *Sulfolobus* Type III-B modules. Cmr-b exhibits RNAse activity cleaving at U-A dinucleotide pairs (Zhang et al. 2012) whereas most other Type III-B complexes cleave at regular spatial intervals along the target RNA, via their Cmr4 subunit (Staals et al. 2013; Benda et al. 2014; Hale et al. 2014).

Cluster 42, like cluster 5, contains an uncharacterised Type III-D core gene encoding the Cmr5 small subunit analog of a subclass of Type III systems which, to date, are poorly characterised. The genes are always located within the Type III-D gene cassette and are co-transcribed with the other Type III-D interference genes (Figure 1E). The subclass of Type III-D modules carrying the gene cluster occur in the bacterial species *Campylobacter, Helicobacter, Fibrobacter, Tannerella* and *Saprospira.*

### Least significantly associated gene clusters

Some gene families were enriched adjacent to Type III genetic modules but were predicted not to be cofunctional. The ten lowest ranking Type III-associated gene families according to the Cas association score are summarised (Table 3). They are dominated by different transposase domains, and include a single ABC transporter family, all of which are ubiquitous in the mobilome and, therefore, less significant. The result reinforces that Type III genetic modules tend to lie in genomic regions with relatively high HGT activity. The result also underlines the importance of employing criteria for coevolution and specificity, in addition to conservation, when searching for *cas* accessory genes.

**Table 3:** The ten lowest ranking genes in terms of significance of association to Type III modules, based on the Type III association score. Even though these gene clusters are often found near genomic Type III systems, they are inferred not to bear any functional link with them, and, therefore, were not considered accessory.

| cluster | size | score | example locus | domain matches | comments |
| --- | --- | --- | --- | --- | --- |
| 209 | 3 | -88.9 | CFBS_2969 | InsA | putative transposase |
| 82 | 6 | -72.2 | UDA_2812 | rve / Tra5 | integrase/transposase |
| 144 | 4 | -66.3 | Pcal_0278 | COG5552/DUF2277 | unknown function |
| 12 | 39 | -60.4 | SSO1518 | Tnp 1 / InsE | transposase |
| 54 | 8 | -46.7 | MAF_28180 | Int C / XerC | tyrosine recombinase |
| 48 | 9 | -42.6 | YN1551_2375 | InsG / IS 4 | transposase |
| 169 | 3 | -33.3 | Cagg_3808 | Ftn | nonheme ferritin |
| 8 | 51 | -33.0 | Smar_0302 | PotA / MalK | ABC transporter |
| 153 | 3 | -32.6 | Athe_0143 | AmyAc MTase | alpha amylase |
| 154 | 3 | -30.33 | Athe_0142 | Yqil | trans-membrane |

### Diversity of accessory proteins with no CARF domain

Although CARF proteins are the most common accessory proteins, their diversity is quite limited with the CARF domain linked to different combinations of a few domains including HTH, WYL, HEPN and PIN toxins. In contrast, the non-CARF accessory proteins appear more diverse, exhibiting a wider range of protein domains and their genes span many smaller gene clusters. Nevertheless, some domain classes are more common than others and they are covered below.

Nucleases constitute the broadest class of non-CARF accessory proteins, associated with clusters 37, 45, 58, 139, 166, 187 and 212. Clusters 37 and 45 encode unrelated restriction endonuclease domains and cluster 37 is probably a core *cas* gene family for the *Sulfolobus* subtype III_V_-1 system, judging from observed gene syntenies. The protein may associate with the Type III effector complex, and facilitate DNA targeting. Cluster 45 genes may encode restriction activity against invader DNA and complement Type III DNA targeting activity. Cluster 166 proteins give good matches to the Nob1 rRNA maturation endonuclease and may be involved in crRNA maturation because the associated Type III modules are not linked to a *cas*6 gene. Nucleases encoded by clusters 58, 139 and 187 are all associated with helicase domains, either as a separate domain in the protein or encoded by co-transcribed genes. Type III systems lack the processive invader dsDNA digestion characteristic of Type I systems, via Cas3, and helicase/nuclease combinations such as the above may contribute the same type of functionality to Type III systems.

Proteases and peptidases represent an important class of non-CARF accessory proteins encoded by clusters 11, 43, 47 and 206. Clusters 43 and 206 encode relatively long proteins possibly with multiple domains that remain of unknown function. Clusters 11 and 47 encode small single domain proteins (about 100 aa), most of which match aspartic acid peptidases. Some cluster 11 members carry no identifiable domains, reminiscent of core SS proteins and may have co-clustered with the peptidases as a result of sequence similarity cut-offs being set too low. However, genes encoding viable peptidases are often found within operons of the cognate Type III Cas proteins which suggests that they are core effector genes and may be involved in maturation of Cas proteins.

ABC and AAA ATPases comprise another well represented group of putative non-CARF accessory proteins with cluster 17-encoded SMC ATPase the most prominent (Figure 1B). Clusters 80 and 93 also encode ATPase domains. The former produces a large protein matching DUF499 in Pfam that probably contains multiple domains and resembles SMC and RecF ATPases at the C terminal end. Cluster 93 encodes a small protein and often accompanies cluster 80, with their genes sometimes co-transcribed, and they carry an AAA ATPase motif. Cluster 35 encodes a protein family found mainly in cyanobacteria; it has a Csx3 domain in the N-terminal end and an AAA ATPase domain in the C-terminal region. Cluster 67 is similar but encodes domains organised in the opposite order. Csx3 is predicted to be a distant member of the CARF superfamily (Topuzlu and Lawrence 2016), and the AAA ATPase is likely to be under allosteric regulation from the CARF domain, with a cOA signal as possible activator.

Another class of accessory gene families are linked to Type III adaptation modules. Members are often located adjacent to adaptation genes including *cas*1 and *cas*2. Cluster 227 encodes a primase domain and is found with adaptation modules of Type III-A and Type III-B systems. A related gene occurs adjacent to Type III-D-linked adaptation modules (Figure 1E). Functions may include DNA repair following a spacer acquisition, thus replacing Cas4, or reverse transcription to generate DNA spacers from RNA invaders. The cyanobacterial Type III-B associated clusters 216 and 27 (Figure 1A) encode small proteins and the former matches a part of Cas1 from Type III-A-associated adaptation modules of other bacteria. Since Type III interference is protospacer adjacent motif (PAM)-independent, this Cas1-associated domain may have evolved to bypass the requirement for a spacer acquisition motif (SAM) (Shah et al. 2013) that occurs in Type I and II systems.

A final class of putative accessory gene families includes those encoding proteins with no significant domain matches. Most of them are small (100 to 200 aa) but a few are larger including those of cluster 124 (average 430 aa). Gene families without identifiable encoded domains are sometimes associated with other accessory genes, including the Cas_RecF-associated gene families (clusters 83, 84 and 108) (Figure 1B) (Vestergaard et al. 2014), the CorA-associated cluster 96 (Figure 1C) and cluster 38 which is often associated with cluster 43-encoded proteases. Other clusters occur alone (cluster 76) or show no clear association pattern (cluster 69), indicating that they can function independently from other accessory proteins.

Next we present two experimental sections to gain further insights into the potential roles and significance of a few accessory proteins. The first constitutes a study of the cyanobacterium *Synechocystis* sp. PCC 6803 where a series of accessory genes were selectively deleted and the effect on Type III-B interference was investigated. In the second, the dependence of RNAse activity on cyclic adenylates was examined for accessory proteins of Type III-B systems of *Sulfolobus* with a view to compare the results with those obtained earlier on bacterial Type III-A systems.

### Type III-B interference activity was not directly impaired in candidate accessory gene deletion mutants

The major defense plasmid pSYSA, present in the cyanobacterium *Synechocystis* sp. PCC 6803, harbours three separate CRISPR-Cas systems - CRISPR1, CRISPR2 and CRISPR3. CRISPR1 and 2 are classified as subtype I-D and III-D CRISPR-Cas systems, respectively (Hein et al. 2013; Scholz et al. 2013; Reimann et al. 2017), while CRISPR3 is a subtype III-B variant system (III-Bv) carrying an unusual fusion of Cmr1-Cmr6 and lacks an obvious Cas6 homolog (Behler et al. 2018) (Figure 2). Recently, it was demonstrated that CRISPR3 is a viable CRISPR-Cas system that functions independently of the non-cognate Cas6 endonucleases associated with CRISPR1 and CRISPR2 (Behler et al. 2018). In addition, the host-encoded RNase E was shown to perform crRNA processing in *Synechocystis* sp. PCC 6803 (Behler et al. 2018).

**Figure 2:**
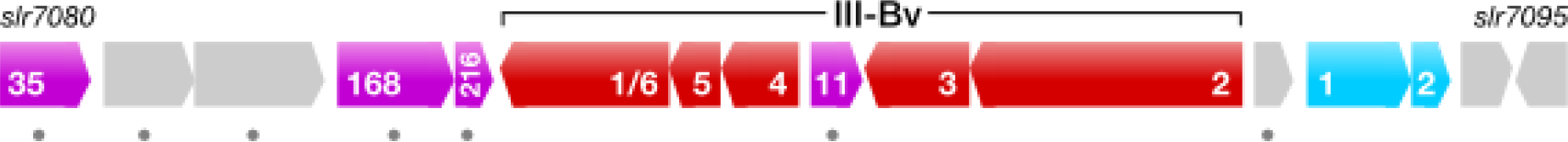
Gene map of the *Synechocystis* sp. PCC 6803 pSYSA Type III-Bv module with flanking genes. Core Type III genes are coloured red and denoted with Cmr numbers. Adaptation module genes are coloured blue and marked with Cas protein numbers. Accessory genes found in this study are coloured purple and indicated with cluster numbers. Genes deleted in mutants subject to the interference assay are marked by a dot below. None of the individual deletions resulted in a marked decrease in interference activity.

To test the functionality of the CRISPR3 system in wild type *Synechocystis* sp. PCC 6803, an interference assay was developed based on two invader plasmids, each containing the reporter gene gentamicin and a fused protospacer sequence in either sense or antisense orientation (Behler et al. 2018). In this study, the same assay was used to test the effects of single candidate accessory gene knock-outs on interference activity. For each accessory gene knock-out mutant, a significant reduction in the number of transconjugants was observed for both invader plasmids relative to the control (Figure 3); this was also observed previously for wild type *Synechocystis* sp. PCC 6803 (Behler et al. 2018). We conclude that the gene products of the investigated accessory genes are not directly involved in the efficient degradation of invading nucleic acids but may participate in other CRISPR-Cas-related processes.

**Figure 3:**
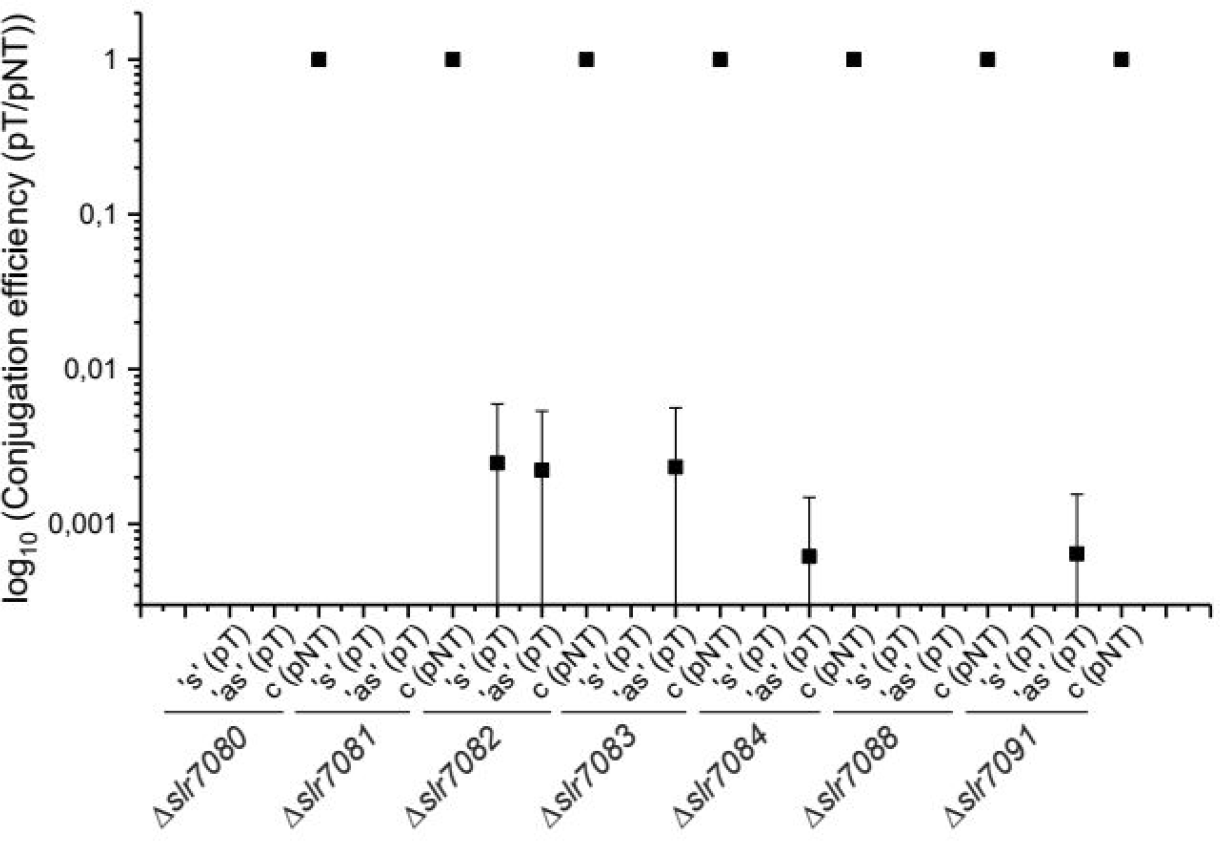
Interference activity of subtype III-Bv-associated accessory gene knock-out mutants in *Synechocystis* sp. PCC 6803. Conjugation efficiencies are calculated by the ratio of the plasmid target (pT) to the plasmid non-target (pNT, control). The conjugation efficiency of the control plasmid was set to 1 and the number of colonies for the plasmid targets was normalized to the control plasmid. Data points represent mean values and standard deviations were calculated for three independent biological replicates. Data points with 0 values are not shown because log(0) is not defined. ‘s’, invader plasmid with protospacer in sense orientation, ‘as’, in antisense orientation, ‘c’, control plasmid without protospacer.

## Compatibility between SisCmr-α and different Sulfolobus CARF proteins

Most putative accessory gene families, especially those encoding CARF proteins, can associate with different Type III subtypes (Table 2). Moreover, highly similar CARF genes, including *sisCsx1* (locus tag: SiRe_0884) are also located adjacent to gene cassettes of different Type III subtypes (Figure 4). This suggests that some degree of compatibility can occur between CARF proteins and different Type III subtypes.

**Figure 4:**
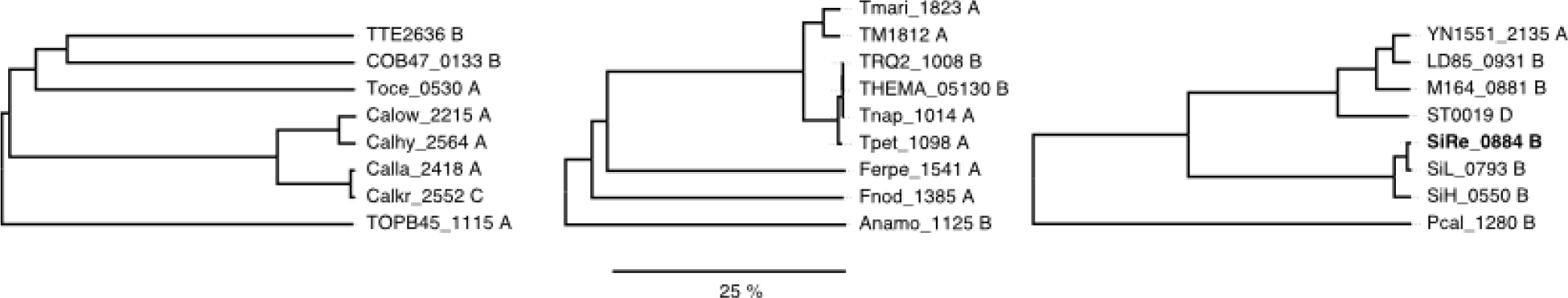
Three subtrees from a neighbor-joining tree of all CARF proteins found in this study. Gene (locus) ids are shown along with the subtype of the associated Type III system. The branch length corresponding to a 25 % dissimilarity at the amino acid sequence level is indicated with the ruler. Closely similar CARF proteins can associate with, and cofunction with, different subtypes of Type III systems. SisCsx1 is included in the final subtree (in bold).

The Type III-B system (Cmr-α) of *Sulfolobus islandicus* (Sis) synthesises cyclic tetra-adenylate (c4A) which binds to the CARF domain of SisCsx1, the cognate accessory RNase of Cmr-α, producing strong RNase activity (WH, QS unpublished). We examined the potential for c4A to activate three additional *Sulfolobus* CARF proteins, namely SiRe_0765 (Csa3), SiRe_0811 (Csm6) and ST0035 (a CARF-PIN toxin) from *S. tokodaii*. ST0035 was shown, by UV crosslinking, to weakly interact with c4A but it did not induce additional RNAse activity. SisCsx1 was the only protein that interacted strongly with c4A, leading to efficient induction of RNA cleavage by activation of the HEPN toxin domain. It was concluded that the other *Sulfolobus* CARF proteins may be activated by other species of cOAs synthesised by different Type III effector systems within the cells.

Although the comparative genomics results suggest that compatibility can occur between CARF proteins and different Type III subtypes, the experiment indicates that this is not a general rule and that other factors may be important for the CARF protein-Type III association to be productive.

## Discussion

The computational method developed for identifying putative accessory genes employed similarity cut-offs and criteria for defining clusters based on considerable experience with manual annotation of genomic CRISPR-Cas cassettes. The Type III association score, based on criteria for host genome self-similarity, Type III specificity and coevolution, yields good granular control over sensitivity versus specificity. The cut-off employed involves a tradeoff which produced a few false positives and some false negatives. However, we infer that most of the gene clusters selected (around 95%) comprise true accessory gene families (Table 2). The method relies on accurate prior annotation of Type III core genes so that flanking genes can be evaluated; for inaccurately annotated Type III systems the method may identify core *cas* genes as accessory genes. Thus, the method will improve with more accurate annotations, and more input genomes, when significantly associated genes will become easier to identify.

The fact that the most widespread CARF and non-CARF accessory genes were identified previously (Table 2) does not undermine the significance of our approach. Accessory protein families have been defined and annotated over a decade involving extensive manual comparisons and curation whereas the present method is semi automatic and scalable and can accelerate the discovery of non-CARF accessory proteins, especially when coupled to an automated pipeline for annotating CRISPR-Cas systems in newly sequenced genomes, an approach that is being developed (Makarova et al. 2015). The present study indicates that the diversity of non-CARF accessory proteins is still under-sampled whereas the major families of CARF accessory proteins have probably been identified (Makarova et al. 2014). Searching additional diverse genomes, and metagenomes, should help identify new accessory protein families with increased precision. Therefore, we plan to integrate the current method into our earlier methods (Lange et al. 2013; Alkhnbashi et al. 2014; Vestergaard et al. 2014; Makarova et al. 2015; Alkhnbashi et al. 2016) and generate a fully automated annotation and characterisation of CRISPR-Cas systems.

### Detection of previously confirmed accessory proteins

Thirty seven of the 76 accessory gene families have been noted in previous comparative genome studies (Table 2). The most important group comprise CARF proteins that were characterised at an early stage (Haft et al. 2005) and were more recently catalogued systematically together with WYL domain proteins (Makarova et al. 2014). The major families of characterised accessory proteins lacking CARF domains include: (a) clusters 5 and 42 (Csx19 and 24) that are annotated as CRISPR-associated in the TIGRFAMs database, and are both likely to encode core Type III SS proteins; (b) clusters 57 and 58 encode the HerA/NurA helicase-nuclease pair and were initially found associated with crenarchaeal thermoneutrophile Type III systems (Bernick et al. 2012) and later were found in other crenarchaea and aigarchaea (Vestergaard et al. 2014); (c) cluster 29 encodes a CorA-like putative magnesium transporter and was originally found linked to Type III-B systems in *Methanococcus* and *Clostridium* (Vestergaard et al. 2014); here we shows it also occurs adjacent to Type III-D systems and in diverse bacteria; (d) cluster 11 or *csx15/20* encodes a Type III-associated peptidase that is erroneously catalogued in CDD as being Type I-U associated (Makarova et al. 2011); (e) cluster 47 encodes a Type III-associated peptidase found together with cluster 37 encoding a nuclease; both are associated with the crenarchaeal III_V_-1 variant subtype (Vestergaard et al. 2014) and may comprise core *cas* genes for this III subtype; (f) cluster 17 or Cas_RecF encodes a repair-associated ABC ATPase found adjacent to diverse crenarchaeal Type III genetic modules (Vestergaard et al. 2014) and it is often accompanied by clusters 83, 84 and 108 which do not yield significant sequence database matches. ATPases, although common and functionally diverse, are well conserved sequence-wise and while a functional link to DNA repair is possible it could be an artefact. However, ATPases could provide the energy required for shifting the chemical equilibrium towards more efficient cleavage of target nucleic acid, given that Type III systems lack the helicases of Type I systems that ensure processive target cleavage.

### Compatibility between CARF accessory proteins and different Type III subtypes

The results (Table 2, Figure 4) suggest that CARF accessory proteins in particular are able to cofunction with various Type III systems regardless of subtype. A mechanistic rationale for this compatibility could be the shared capacity of Type III subtypes to synthesize the cOA signal required to activate CARF proteins upon invader RNA recognition. The CARF domain is likely invariant for Type III modules as long as a cOA messenger can activate it. While this explains the observed subtype compatibility, experimental evidence from Type III-A systems in *Streptococcus thermophilus* (St), *Thermus thermophilus* (Tt) and *Enterococcus italicus* (Ei) (Kazlauskiene et al. 2017; Niewoehner et al. 2017) and a Type III-B system in *Sulfolobus islandicus* (Sis) (WH, QS unpublished), suggest that the mechanisms are more complex (Table 4). The different Type III-A Csm effector complexes tested produced different cOA profiles; some synthesising mostly hexa-adenylates (c6A) while others produced mainly triadenylates (c3A). The archaeal Type III-B system synthesised a tetra-adenylate (c4A) signal (WH, QS unpublished). Moreover, the synthesis efficiencies varied by more than an order of magnitude between the different Csm and Cmr effector complexes tested (Table 4) and the *E. italicus* system had to be genetically modified to produce significant yields. In addition, the CARF RNAses tested showed strong preferences for specific cOA species, with some favouring c6A and others c4A and, moreover, their activation efficiencies varied markedly, with some CARF proteins requiring several orders of magnitude higher cOA concentrations to trigger efficient RNA digestion.

**Table 4:** Comparisons of reaction efficiencies of different CARF proteins with respect to RNAse activity, and for Type III effector complexes with respect to cOA synthesis. Substrates comprise RNA for CARF proteins and ATP for Type III complexes. Reaction times, substrate and effector concentrations shown are the minimum required for digestion of 50% of the RNA substrate or for converting more than 80% of ATP into cOA. The concentration of cOA required for activation of CARF proteins differs by several orders of magnitude, as does the efficiency with which the Type III complexes synthesize cOA. The c6A required to activate StCsm’s cognate Csm6 protein comprises a minor species (only 0.5% total cOA synthesised), with the major species being c3A. In contrast, almost all cOA synthesised by SisCmr and EiCmr was of the type required by cognate CARF proteins. Table contents were compiled from published data (Kazlauskiene et al. 2017; Niewoehner et al. 2017) while the SisCsx1 and SisCmr data were produced in the Copenhagen laboratory (WH, QS unpublished).

|  | SisCsx1 | TtCsm6 | StCsm6 | StCsm6' | EiCsm6 | SisCmr | StCsm | EiCsm |
| --- | --- | --- | --- | --- | --- | --- | --- | --- |
| class | Csx1 | Csm6 | Csm6 | Csm6 | Csm6 | III-B | III-A | III-A |
| active cOA species | c4A | c4A | c6A | c6A | c6A | c4A | c6A | c6A |
| effector conc (nM) | 100 | 10 | 0.1 | 1 | 0.15 | 10 | 200 | 160 |
| Substrate | 520 nM | 10 nM | 10 nM | 10 nM | 40 nM | 100 μM | 50 μM | 500 μM |
| cOA (nM) | 20 | 500 | 0.5 | 5 | 5 | ~ 100 % | 0.5 % | ~ 100 % |
| reaction time (min) | 20 | 30 | 30 | 30 | 4 | 20 | 10 | 30 |

The cOA specificities of CARF proteins and the effector complexes appear to transcend the broad protein family, and effector subtype categories, because they presumably vary over shorter evolutionary distances. For example, the Type III subtypes are not linked to specific cOA species (Table 4) and such mechanistic differences probably occur at a finer subfamily level.

CARF accessory proteins and core Type III effectors must be able to cofunction in order for their linked genes to persist. However, the experiments summarised above indicate that variables such as the sensitivity to the signal, the species and its concentration, limit productive CARF-Type III interactions.

### Mobilome-associated genes adjacent to Type III modules

In addition to accessory gene families, we expected mobilome-associated gene families to be enriched around genomic Type III *cas* modules. The Type III association score was devised to address this problem, and five out of the ten lowest ranking gene clusters encoded putative transposases (Table 3). Remarkably, no toxin-antitoxin (TA) gene pairs were found despite (a) previous studies showing that they are common in mobilomes, with some bordering CRISPR-Cas modules (Guo et al. 2011; You et al. 2011) and (b) additional evidence demonstrating coevolution between TA gene pairs and adjacent Type III modules (Shah and Garrett 2013). The TA systems may influence mobility of adjacent CRISPR-Cas modules or they may help induce dormancy or programmed cell death at key stages of the immune response, in order to minimise the spread of invader genetic material in the cellular population (Makarova et al. 2012). Thus, they could provide a semi-accessory function to Type III immunity while also being able to function independently. This would explain why some specific subfamilies of TA systems are enriched adjacent to CRISPR-Cas gene cassettes while others are not (Shah and Garrett 2013) and, as a result, they were not detected by our method.

Of the many accessory proteins identified that yield no protein domain matches with databases some of these could be anti-anti-CRISPR proteins. An increasing number of anti-CRISPR proteins have been characterised, encoded by phages (Borges et al. 2017) and by archaeal viruses (He et al. 2018), and it remains a possibility that some of the CRISPR-Cas systems encode accessory proteins that can neutralise the anti-CRISPR proteins before they inhibit an immune response.

### On the role of Type I associated CARF proteins

Type III effectors activate CARF proteins via a cOA messenger molecule signaling the presence of intracellular invader RNA (Kazlauskiene et al. 2017; Niewoehner et al. 2017; Han et al. 2017b) and many CARF protein genes were identified earlier adjacent to Type III modules (Makarova et al. 2002; Haft et al. 2005). Less clear is why CARF protein genes are common adjacent to Type I systems, especially amongst archaeal hyperthermophiles (Vestergaard et al. 2014), when Type I core Cas proteins do not apparently possess the GGDD domain responsible for cOA derivatives. Most types of CARF protein genes that are associated with Type III modules, are occasionally found adjacent to Type I cassettes, with *csx3* and especially *csa3* being the most common. *csa3* encodes a transcriptional repressor of Type I-A interference modules with a DNA binding HTH domain coupled to a ligand sensing CARF domain. Its pattern of invariant presence suggests that it is a core gene for that subtype.

Type I-A gene cassettes in several crenarchaeal thermoacidophiles, including species of *Sulfolobus, Thermosphaera, Thermogladius, Fervidicoccus* and *Desulfurococcus* carry two *csa3* genes and recent studies have shown that one encodes a transcriptional repressor of Type I interference (He et al. 2016) while the other activates the adaptation module (Liu et al. 2017). The former study detected invader nucleic acid-dependent activation of transcription of the Type I-A interference gene cassette. Moreover, a model was proposed that involved Csa3 binding to the Type I-A effector complex which then bound to the interference module promoter and repressed further transcription. Protospacer-containing DNA then recruited the effector complex and derepressed the Type I-A interference operon. A more likely model, given our current knowledge of Type III systems involves sensing of invader nucleic acid by a Type III effector complex and, after recognition of invader RNA, the cOA signal produced may derepress the Type I-A interference module by inducing a conformational change in its CARF domain-containing transcriptional repressor. This model would also imply that spacer acquisition is under the positive control of Type III-mediated invader sensing, reminiscent of the primed adaptation seen for other Type I subtypes but involving a different mechanism. Thus all CARF proteins associated with Type I systems may facilitate cofunctioning with Type III systems, either directly, as for Csa3, or indirectly by selecting for two types of CRISPR-Cas systems. Genomes that harbour Type I systems flanked by CARF genes like *csx1* are thus more likely to exhibit and maintain Type III systems, even when located on a distant genomic locus. This model also implies that Type I-A systems can be dependent on Type III systems for activating spacer acquisition an/or effector functions. Such an interdependence could give rise to synergies that would ensure a more robust immune response in a natural environment with extensive viral diversity, especially as occurs amongst archaeal hyperthermophiles.

### Type III systems as a general purpose RNA interference platform

Our finding that deletion mutants of *Synechocystis* sp. PCC 6803 Type III-Bv accessory gene candidates were not deficient in interference activity confirmed that RNA silencing and transcription-dependent DNA silencing are inherent to core Type III functionality. The additional functions provided by the deleted accessory genes are likely to lie beyond basic interference.

The extensive diversity of non-CARF accessory gene families located adjacent to Type III interference modules, suggests that they are multifunctional, likely extending functionality beyond immune defence. The basic RNA interference activity of core Type III effectors is non-processive in contrast to Type I interference activity where Cas3 processively digests invader DNA after protospacer recognition (Sinkunas et al. 2011; Beloglazova et al. 2011; Westra et al. 2012). Type I systems are streamlined towards efficient invader dsDNA degradation, whereas core Type III systems are more versatile interfering with different types and configurations of nucleic acid. Thus Type III systems may be more useful for invader surveillance than clearance, and the cOA signaling pathway is ideal for directing effector functionality to other cellular systems. The specialisation of Type I systems towards supercoiled dsDNA targets suggests that they could have evolved from a more general purpose Type III-like ancestor, after the addition of a helicase-nuclease pair. This is also consistent with a recent hypothesis for evolution of Type I systems from Type III systems (Koonin and Makarova 2017). Adaptive immunity, although important, could well be one specialisation from a pool of functions offered by Type III systems. RNA interference has a variety of potential applications beyond immunity, particularly within information processing, and different accessory proteins may provide the crucial functional links to other cellular systems.

### Concluding remarks

The developed computational method successfully identified 76 putative accessory gene families flanking genomic Type III genetic modules of archaea and bacteria, more than half of which had not been identified previously. The results expand in particular the repertoire of diverse non-CARF accessory gene families for which functions are currently unknown. The diversity of the gene families found, provides evidence of Type III systems being coupled to numerous functions additional to invader nucleic acid silencing. The study also suggests that more accessory gene families will be detected in the future when automated annotation of an increasing number of genomic CRISPR-Cas cassettes becomes available.

## Materials and Methods

The comparative genomics method for detecting putative accessory genes employs a “guilt-by-association” approach, where genes flanking known Type III modules are first clustered by sequence similarity. The extent of conservation across a wide range of host genomes is taken as an indication that the flanking gene clusters are functionally linked to their cognate Type III modules. Since genomic CRISPR-Cas cassettes are most often located in genomic regions implicated in regular horizontal gene transfer (HGT), the mobilome, any genes flanking the cassettes will be shuffled even over short evolutionary distances. The method assumes that gene conservation adjoining Type III gene modules over larger evolutionary distances stems from a selective pressure to maintain those genes in close vicinity of the cognate Type III systems, indicative of a likely functional link. The mechanistic rationale underlying this, is that any horizontal transfer event involving a Type III module will be more likely to include any co-functional accessory genes the closer they are located to the core genetic module. This would ensure the transfer of fully functional cassettes into a new host and provide it with a selective advantage, and increase the chances of the cassette replicating in the new host.

The method used for identifying putative accessory genes relies on prior accurate annotation of core Type III genes. Therefore, the data set of annotated Type III modules from the latest CRISPR-Cas classification update is used (Makarova et al. 2015). For less well annotated Type III modules, the method is expected to pick up core genes in addition to any accessory genes. Furthermore, some gene families with no functional link may border Type III genetic modules and be enriched. Such gene families may include transposases, toxin-antitoxin gene pairs and genes within mobile genetic elements, all of which are generally enriched in mobilome regions. To distinguish true accessory genes from such false positive gene clusters, the Type III specificity and degree of Type III coevolution is estimated for each gene cluster. Gene clusters that have a history of accompanying Type III systems are predicted to display a high degree of coevolution and specificity for their cognate Type III module, and the results can thus be used to remove spurious gene clusters from those that are functionally conserved. By setting optimal cutoffs for defining gene clusters, specificities and estimates for coevolution, it is expected that most genes that pass them comprise accessory genes. While false positives may still occur, the chances of them being selected is minimised.

### Definition of putative accessory gene clusters

Five genes upstream and downstream of annotated Type III gene modules were selected from the most recently classified archaeal and bacterial CRISPR-Cas systems (Makarova et al. 2015). They were then pooled and subjected to an all-against-all sequence similarity comparison and the genes were then clustered according to their protein sequence similarities. Sequence clusters were defined to be as inclusive as possible without unrelated protein sequences falling in the same cluster. This was achieved by optimising the sequence similarity cut-off and, also, by defining a custom sequence similarity metric that took into account total protein sizes in relation to alignment length and the relative locations of similar sequence regions between the two compared proteins. The custom similarity score was computed for each protein pair and forwarded to Markov clustering (Enright et al. 2002) which generated the clusters. A multiple sequence alignment (MSA) was made for each gene cluster and used for sensitive profile-profile sequence searches (Söding et al. 2005) against the PFAM, CDD, TIGRFAMs and COG databases (Punta et al. 2012; Makarova et al. 2011; Haft et al. 2005; Makarova et al. 2006) as well as the latest version of the PDB70 database that is distributed together with the HHSuite package (Söding et al. 2005; Burley et al. 2017). MSAs were also used for a phylogenetic analyses in order to clarify whether the gene clusters had co-evolved with their cognate Type III systems. Subclusters were defined for each cluster in order to estimate diversity and as a fall-back option when the main clusters were too broad. Hidden Markov models (HMMs) corresponding to each cluster were constructed and used for genome-wide searches to quantify the specificity of each gene cluster with regard to Type III gene module association. Pairwise sequence similarity searches with the individual cluster member sequences as queries were also used for this purpose. A *cas* association score was then devised to distinguish spuriously associated gene families from those that were significantly associated. The score combined the coevolution data with data on genome-wide Type III specificity, along with information about the genomic similarity of hosts carrying the gene cluster. Thus a gene cluster seen conserved across a limited range of closely related host genomes was downgraded compared to a gene cluster conserved adjacent to type III systems across widely divergent hosts. The *cas* association score was used to rank all conserved gene families, and only the top-ranking gene families were included for further analyses, with lower ranking genes families assumed to be spuriously associated. Gene clusters corresponding to previously confirmed Type III accessory genes (Vestergaard et al. 2014; Makarova et al. 2014) we included regardless of association score. Graphical representations of the gene neighbourhoods surrounding all surveyed type III modules were then created and significant accessory genes were marked in order to inspect the results visually and individually.

### *Synechocystis* interference assay

The self-replicating conjugative vector pVZ322 and the gentamicin resistance cassette were used for the construction of invader plasmids to check interference activities in *Synechocystis* sp. PCC 6803 as described in (Behler et al. 2018). Spacer 2 was selected to derive an appropriate protospacer sequence and the protospacer sequence was fused in frame to the gentamicin resistance (reporter) gene just upstream of its stop codon in both orientations, sense and antisense, respectively. Mean values of conjugation efficiency and standard deviations were calculated by the ratio of the number of conjugants of target plasmids to non-target plasmids in biological triplicates and in parallel for the control plasmid. Interference activity was observed if the conjugation efficiency was <1. Conjugation efficiency of the non-target plasmid was set to 1, which corresponds to no interference activity.

### Creation of knock-out mutants and transformation of *Synechocystis* 6803

*Synechocystis* sp. PCC 6803 cultures were grown as described (Hein et al. 2013). Single gene knock-out mutants of slr7080-slr7084, slr7088 and slr7091 were generated by integrating a kanamycin resistance cassette into the corresponding loci. The up- and downstream flanking regions (1,000 base pairs in size each) were amplified via PCR to ensure site-directed integration via homologous recombination. The resistance cassette was placed in between the upstream and downstream flanking regions in vector pUC19 by Gibson assembly. Transformation of 10 mL *Synechocystis* sp. PCC 6803 aliquots was performed as described (Hein et al. 2013). Transformants were tested for full segregation of the introduced knock-out via PCR screening.

### *Synechocystis* 6803 conjugation

Plasmids were conjugated into *Synechocystis* sp. PCC 6803 by triparental mating as described in (Scholz et al. 2013) and few variations described in (Behler et al. 2018). Briefly, the helper strain *E. coli* J53/RP4 and the donor strain *E. coli* DH5α with the plasmid of interest are combined to allow the transfer of the RP4 plasmid into the plasmid of interest bearing cells. After the addition of the recipient strain *Synechocystis* sp. PCC 6803, the plasmid of interest is transferred from the *E. coli* donor strain to the cyanobacterial recipient strain by conjugational transfer. 40 µL of cell suspension were plated on BG11 agar plates containing 5 µg/mL gentamicin. Conjugants were counted after further incubation at 30°C for 2 weeks.

### Interaction between c4A and *Sulfolobus* CARF proteins

c4A was synthesized by incubation of Cmr-alpha with ATP and target RNA (WH, QS unpublished). The interaction between c4A and CARF proteins was determined using a UV cross-link assay as described previously (Han et al. 2017b). The extent of activation of the *Sulfolobus* CARF proteins by c4A was determined by measuring RNAse activity as described previously (Han et al. 2017b).

## Acknowledgements

The authors are grateful to all members of the FOR1680 Consortium for helpful discussions.

## Disclosure statement

The authors declare no conflict of interest.

**Supplementary material:** Yes

**Figures:** Yes - 4 figures

**Tables** Yes -4 Tables

